# Metaproteomics Reveals Functional Rewiring and Metabolic Limits in Synthetic Bacterial Consortia Degrading Complex Waste Oils

**DOI:** 10.64898/2026.02.17.706413

**Authors:** Ali Al Haid, Henri Saddler, Hannah Leflay, El Baraa Elghazy, Kirsty Franklin, Caroline Evans, Jagroop Pandhal

## Abstract

Elucidating the metabolic mechanisms underpinning complex hydrocarbon biodegradation by microbial consortia benefits from integration of community-level genomic and proteomic data. Here, we deployed a combined metagenomics and metaproteomic analysis to characterize functional dynamics within bacterial consortia enriched from hydrocarbon-contaminated soil, using military waste oil as the sole carbon source. Metagenomic profiling revealed a consortium dominated by *Stenotrophomonas maltophilia* alongside established hydrocarbon-degrading taxa including *Pseudomonas fluorescens*, *Pseudomonas putida*, and *Delftia acidovorans*. Quantitative label-free metaproteomics uncovered dynamic taxonomic restructuring and metabolic reorientation, with *Delftia* and *Achromobacter* proteins enriched during active degradation, indicating niche advantages under chronic contamination. Enzymatic mapping resolved expression of degradation pathways targeting aliphatic and aromatic substrates, with one specific consortium, BPS4, exhibiting preferential catabolism of catechol intermediates channeled toward the tricarboxylic acid cycle, supported by elevated protein abundance of oxidoreductases, dehydrogenases, and β-oxidation enzymes. Gene ontology enrichment revealed heightened nitrogen cycling capacity, including upregulation of nitric oxide biosynthesis and nitrate reductase, suggesting adaptive nitrogen metabolism coupled to hydrocarbon mineralization. *In-vitro* batch-fed adaptation studies demonstrated limited improvements in degradation of recalcitrant fractions, particularly branched aliphatics and organophosphates, revealing intrinsic metabolic constraints within the consortium. These findings underscore the complementary utility of metaproteomic interrogation for resolving both population-level dynamics and functional metabolic capabilities, while highlighting fundamental limitations in expanding the degradative spectrum of enriched consortia through conventional adaptation strategies. This work establishes a foundation for rational ecological engineering of hydrocarbon-remediating consortia informed by metaproteomics.

**Importance:** Waste oils from transport and industry are complex and toxic mixtures that are difficult and expensive to treat using conventional chemical methods. Microorganisms offer a sustainable alternative, but their performance depends on how different species work together and where their metabolic limits lie. In this study, we show how advanced protein-based analysis using quantitative metaproteomics, can reveal microbial community activities whilst breaking down real-world waste oils. We enriched bacteria from contaminated soils and then identified the enzymes driving effective degradation. We investigated how communities reorganise under chemical stress, and why certain oil components remain resistant even after long-term adaptation. By uncovering both strengths and limitations of naturally enriched microbial consortia, this work provides practical insights for designing more reliable biological treatments for oil-based wastes and even contaminated environments, supporting greener approaches to pollution management and industrial waste recycling.

## Introduction

Petroleum contamination of soils and aquatic systems has multiple devastating environmental impacts, including disruption of ecological food chains through bioaccumulation of toxic components (1). It is widely accepted that majority of global oil pollution comes from anthropogenic activities, particularly from the petrochemical industry. The Exxon Valdez tanker spill in 1989 at Prince William Sound, Alaska, resulted in 40.8 million liters of crude oil causing significant mortality in wildlife, in addition to subsurface oil layers persisting in sediments for >20 years (2). Similarly, the tragic incident of the Deepwater Horizon on the Gulf of Mexico USA in 2010 caused by a blowout during offshore drilling due to faulty cementing and pressure management failures, resulted with 206 million gallons of crude oil released creating contaminated shorelines of up to 800 km, in addition to long term consequences on the biodiversity of coral reefs (3, 4). Though these are globally known examples of mega oil pollution incidents, petrochemical ecotoxicology is also caused by the exponential growth of modern global transportation and industries such as motor vehicles, aviation, and maritime shipping. Global oil waste streams can be divided into several categories, where waste motor oils (WMO) and waste industrial oils (WIO) are used as raw materials for regeneration and purification, while waste oil mixtures (WOM) are processed at the refinery in a mixture with crude oil. Global accumulation of WOM is ∼15 million tons/year, and in Europe alone up to 1.6 million tons/year is either collected or disposed of, where collected oils are either regenerated, used as fuel, or destroyed (5). The regeneration of waste oils presents overall economic advantages by recycling and repurposing waste oils, while contributing towards less adversity towards the environment. Recycling of waste oils follows a set pipeline that initiates by removal of impurities, followed by dehydration, evaporation of low-boiling point hydrocarbons, removal of hydrocarbon transformation products, and supplementation of additives (5). Physicochemical and thermochemical processing methods have been extensively developed to mitigate environmental concerns and comply with legislative restrictions associated with conventional recycling approaches. As an alternative to recycling, biological treatment or breakdown of waste oils is proposed to as a more ecologically friendly strategy, however biodegradation pipelines have yet to be cost-effectively implemented within large-scale industrial operations.

It has been widely regarded that biological treatment of hydrocarbons can have major environmental and economic advantages, though it still faces techno-economical obstacles when scaling up from laboratory models. Over the decades, research on notable microorganisms capable of degrading petroleum hydrocarbons have focussed on bacteria due to their wide and versatile application using a diverse range of enzymes (6). Despite the increased number of identified genera and species of hydrocarbon degrading bacteria (HDB), no single bacterial strain is capable of complete mineralization of WOMs alone, especially mixtures consisting of complex and heavy hydrocarbons (6). This has led to the prospect of using microbial consortia for complete mineralisation of hydrocarbon mixtures, where specific hydrocarbons and by-products are targeted by different bacteria. Enrichment of consortia for this purpose is undertaken from environments contaminated with petroleum hydrocarbons, which is paramount for isolation of key HDB genera. This is demonstrated with recent studies capable of isolating species and consortia with high toxicity resistance from traditionally difficult fractions of petroleum, such as heavy oil, oil sludge, and asphaltenes (7–9). Constructing synthetic or engineered consortia for bioremediation can be undertaken through bottom-up or top-down designs. In bottom up approaches, isolated microbial strains are mixed in combinations and screened for degradation capabilities (10). These strains can be further modified using synthetic biology to enhance stability or control. Top-down methods to construct functional consortia is less laborious and does not rely on strain isolation techniques. Therefore, top-down community enrichments better preserve intricate metabolic networks that are indigenous to the microbial members in cohabitation. Though top-down enrichment of microbial communities remains challenging, especially for unculturable microorganisms, studies demonstrated top-down enrichment’s efficacy, stability, and resilience in bioprocessing pipelines such as anaerobic lignocellulose bioconversion, waste water treatment, and oil sludge treatment (11–13). Furthermore, using adaptive evolution to further improve efficiency in top-down enriched consortia has been demonstrated for bioremediation of leachates derived from landfill (14).

To understand the complex interaction mechanisms occurring in synthetic microbial consortia, high throughput molecular analyses technologies, commonly known as “Omics”, have facilitated rapid advancements arguably due to the increased interest in microbiome research. The complex relationship of community members, whether competitive or cooperative, is currently more accessible owing to streamlining and standardization of high-throughput data technologies (i.e. metagenomics, metatranscriptomics, and metaproteomics) driven by optimization and increases of computational power (15–17). Expression profiles can also reveal niche activities, as shown in a metatranscriptomic study of activated sludge systems (18). Ammonia oxidizing and nitrite oxidizing bacteria representing <0.15% and <0.25%, respectively, of the activated sludge community were found to be significantly associated with governing heavy oil biodegradation (18). Similarly, research on polyethylene-associated biofilm communities indicated plastic-degrading gene expression patterns associated with biofilm development, where *alkB1/alkM* transcripts encoding polyethylene biodegrading enzymes were up-regulated during biofilm maturation, along with transporters and fatty acid β-oxidation pathways (19). Metaproteomics can also identify statistically significant changes in community composition in response to sunlight, pollution, and chemical surfactants. Researchers examining surface waters collected off Pensacola Beach Florida, Gulf of Mexico, relatively quantified showed significant alternations in proteins involved in carbon metabolism pathways, photosynthetic protein expression, and nitrogen and phosphorus cycling proteins (20). Moreover, the occurrence of community change can be exploited to guide and develop synthetic bacterial consortia designs using metaproteomics. Macchi et al. (2021) evaluated synthetic bacterial consortia for optimized phenanthrene degradation guided by integrated genomics and shotgun proteomics. By identifying 1,579-1,622 proteins across two communities, functional redundancy and competition dynamics between members was revealed (21). This study reports the construction of two bacterial consortia, enriched from soil obtained from a military site which is polluted with complex WOMs. Bacterial consortia were enriched using WOMs as the sole carbon source supplement, in tandem with characterization of hydrocarbon composition. Enriched bacterial consortia were taxonomically and functionally profiled using a bottom-up metaproteomics workflow, elucidating species and metabolic pathways key in driving waste oil biodegradation. In addition, this work presents the novel application of metaproteomics for artificial selection studies, providing insights into the molecular mechanisms underlying bacterial community adaptation to nutritional and hydrocarbon stress.

## Materials and Methods

### Waste Oils sample

Waste oils (WO) were provided by the Defense and Security Accelerator (DASA) from the Royal Air Force (RAF) Brize Norton Airbase, which were used as substrates for the enrichment of bacteria. The WOs contained undisclosed ratios of a mixture of oils, such as synthetic hydrocarbon “non-petroleum base” aircraft (OX19), Turbine fuel aviation (Jet-A1 FSII), and aircraft lubricating oils (BP Turbo Oil 2380).

### Consortia Enrichment

Soil samples were collected from the RAF Brize Norton airbase at three different sites previously exposed to hydrocarbon contaminants (51°45′00″N 001°35′01″W / 51.75000°N 1.58361°W). Each site of soil collected were annotated as BPS2, BPS4, and BPS8. Soil sites BPS2 and BPS4 were both of clayish consistency in contrast to site BPS8, which was relatively drier. In the case of BPS8, the site had previously undergone remediation for an F34 aviation fuel spill. All samples collected from the sites were between 5-10 cm below the surface, while separating the organic layer, and removing any geochemical matter.

Four types of broth mediums were used for the enrichment process, M9 minimal salts (M9) (5X powder, Sigma-Aldrich, St. Louis, MO, USA); Reasoner’s 2A (R2A) (22); Mineral Salts Media (MSM) (23, 24); and Bushnell Haas media (BH) (25). Each type of media was made up to a set of four 100 mL culture flasks. For each set, two flasks were inoculated with 1g of soil from BPS4 and BPS8 (1% w/v ratio), and one flask as an abiotic control. All flasks were supplemented with 1 g (1% w/v ratio) of the WO mixture as a substrate for enrichment. Cultures were incubated at room temperature (20 °C) at a shaking rate of 120 RPM. Qualitative analysis on physical changes to oil and cultures, including oil miscibility, organic layer fragmentation, culture turbidity and emulsification, were recorded for all flasks within the duration of enrichment.

### Waste Oil Biodegradation Experimental Design

Stock cultures of consortiums BPS4 and BPS8 (3 biological replicates each) were revived in LB Broth and incubated in ambient temperature at 110 RPM until an optical density of 1 at 600 nm wavelength. Each consortium was inoculated at an OD 600nm of 1.0 in a set of Mineral Salts Medium (MSM) cultures. MSM media composition: dipotassium phosphate (K2HPO4) 1.8 g/L, ammonium chloride (NH4Cl) 4.0 g/L, magnesium sulphate (MgSO4) 0.2 g/L, sodium chloride (NaCl) 2.5 g/L, ferrous sulphate (FeSO4) 0.01 g/L, at pH 7, was the medium of choice for this experiment. Positive control culture sets contained 0.1M of sodium acetate in MSM, while experimental cultures were supplemented with 1% (w/v) WOs, as a sole carbon source. Negative abiotic controls contained only MSM medium with 1% (w/v) WOs. Cultures were incubated at ambient temperature, with shaking at 180 RPM, for 14 days.

### Batch-fed adaptation experiment

The adaptation experiment was conducted to evaluate the adaptation potential of bacterial consortia BPS4 under different carbon source conditions using a batch-fed dilution to extinction method of artificial selection. Three types of media were prepared: Mineral Salts Medium (**MSM**), Acetate + Mineral Salts Medium (**A-MSM**), and M9 Minimal Salts + Trace Elements Medium (**TE-M9**). MSM medium was made up of 1.8g/L K₂HPO₄, 4.0g/L NH₄Cl, 0.2g/L MgSO₄, 0.1g/L NaCl, and 0.01g/L FeCl₃. A-MSM medium was made up by adding sodium acetate carbon source at a final concentration of 0.1M. TE-M9 is initially made up by dissolving 5X M9 minimal salts (Sigma Aldrich M6030) composed of 15 g/L KH_2_PO_4_, 2.5 g/L NaCl, 33.9 g/L Na_2_HPO_4_, and 5 g/L NH_4_Cl, subsequently supplemented with 1mL of each 1000X Corning trace elements A (Corning 25-021-CI), B (Corning 25-022-CI), and C (Corning 25-023-CI) to a final volume of 1L. Stock cultures of bacterial consortiums BPS4 were grown overnight in LB broth medium to an optical density of approximately 1 at 600nm wavelength for inoculation. For each consortium, five experimental conditions were established: Control Acetate (A-UN), Control WOs (ROS-UN), Adapted Acetate (A-AD), Adapted WOs (ROS-AD), and Abiotic WOs (negative control). Control and Adapted Acetate culture flasks consisted of A-MSM, while Control and Adapted WOs culture flasks used MSM supplemented with 1% w/v WOs, were subsequently inoculated with the bacterial consortium. All cultures were incubated at ambient temperature with agitation at 180rpm. Control cultures (UN) were incubated for 14 days with no media refreshment. Adapted cultures (AD) were serially transferred once every 14 days for a total duration of 5 months, where consortia are subcultured into fresh TE-M9 medium supplemented with 0.1M sodium acetate (A-AD) and 1% WO (O-Ad) to maintain consistent selective pressure. Abiotic WOs negative control contained MSM and 1% WOs and were refreshed with TE-M9 medium supplemented with 1% WO.

### Matched Metapeptide Database Library

DNA extraction of enriched bacterial consortia was undertaken using the Invitrogen PureLink® Genomic DNA Mini Kit. Both BPS4 and BPS8 were revived and cultured in LB Broth until an optical density (OD) at 600nm of ∼1. Post incubation, bacterial cells were harvested (1 mL of suspension) and resuspended in PureLink® Genomic Digestion Buffer, in addition to Proteinase K (both supplied with the kit), and mixed by vortex to lyse bacterial cells. Bacterial lysates were then incubated in a water bath at 55 °C with intermediate vortexing. RNase A (supplied with the kit) was added to the lysate, mixed well by vortexing, and incubated at room temperature for 2 minutes. PureLink® Genomic Lysis/Binding Buffer and 96 to 100% concentration range of ethanol was then added to homogenize the lysate. Homogenized lysates were then added into a PureLink® Spin Column in a Collection Tube to bind gDNA, centrifuged at 10,000 × g for 1 minute at room temperature, and decanted solution. Bound gDNA was then washed with Wash Buffer 1 (supplied with the kit), centrifuged at room temperature at 10,000 × g for 1 minute, then repeated with Wash Buffer 2 (supplied with the kit). Bound gDNA was then eluted with 10 mM Tris-HCl, pH 8.0–9.0 and stored at –20°C. gDNA extract quality was analyzed through Qubit and Fragment analyzer. Consortiums BPS4 and BPS8 gDNA extract samples were sequenced using the Illumina Novaseq platform, via paired-end sequencing 2×150 bp over 32 increment reads, with an estimate of ∼160M raw reads per sample. Sequence reads outputs (FASTQ files) were analyzed for quality control using FastQC v0.1, developed by the Babraham Bioinformatics. The following tools mentioned were all used via the Galaxy-P multi-bioinformatics web-based platform (26). Metagenome FASTQ files were inputted into TrimGalore! to excise Illumina adapter reads from sequence data. Sequence read fragments were assembled into contigs via the MEGAHIT assembly tool. The MEGAHIT Assembly quality was assessed using the QUAST software. Assembled contigs are then grouped and binned into their respective taxonomic genomes at a species level using MaxBin v2.0. Binned and unbinned assembly databases were inputted into a variety of gene prediction and annotation tools, Prokka, CAT, and FragGeneScan. Peptide sequences of predicted and functionally annotated genes were taxonomically annotated using the Unipept tool via pept2lca application (27). Optionally, functionally unannotated peptide sequences were inputted into the Unipept pep2prot application to retrieve their respective Uniprot IDs, then subsequently used to taxonomically annotate Uniprot IDs via the Retrieve Uniprot Entries tool (<u>GalaxyP Uniprot tool: URL Link</u>) on Galaxy-P. Metapeptide peptide databases were then inputted into cd-hit to reduce redundancy of peptide sequences.

### Liquid-Liquid Extraction of Residual Waste Oils

Cultures supplemented with 1% (w/v) WOs were decanted into 50 mL falcon tubes and were centrifuged at 5000 x g for 15 minutes to separate the organic layer, in addition to pelleting bacterial consortia. N-Hexane was gently added into centrifuged tubes and gently mixed to dissolve the WOs in the organic, without disturbing the aqueous layer and the bacterial pellet. WOs dissolve in n-hexane were carefully extracted via a needle and syringe and placed into clean 30mL screw cap glass vials. Residual WOs adhered to glass cultures and falcon tubes were dissolved with gentle mixing with n-hexane, then pooled into their respective 30mL screw cap glass vials. Glass vials containing extracted WOs were then placed inside fume hoods overnight to allow evaporation of n-hexane. The remaining aqueous layer inside falcon tubes were decanted gently without disturbing bacterial pellet, then phosphate buffer saline was used to wash the bacterial pellet prior to harvesting and storage for cell lysis. Extracted WOs were dissolved in 30mL n-hexane solvent and were acidified with concentrated HCl. Dissolved WOs were then transferred into a separatory funnel and shaken vigorously before allowing the organic and aqueous phases to separate. The aqueous phase was discarded. The organic phase was passed through a glass funnel containing a glass wool bedding along with sodium sulfate to dehydrate the oil phase and break emulsions. The extracted WOs were placed in a fume hood to allow n-hexane to evaporate. Extracted oil samples were then resuspended in uniform n-hexane solvents prior to total petroleum hydrocarbon (TPH) quantification using the InfraCal 2 ATR-SP (Spectro Scientific, Chelmsford, Massachusetts, USA) by infrared spectrometry and GC-MS. Residual WOs were analyzed using gas chromatography-mass spectrometry (GC-MS) employing an Agilent DB-5MS-UI capillary column (30 m × 0.25 mm, 0.25 μm film thickness). Helium was used as the carrier gas at a constant flow rate of 1.2 mL min⁻¹ and a column head pressure of 9.47 psi. Samples (1.0 μL) were introduced via split injection (100:1 split ratio) with the injection port maintained at 250°C. The temperature program was initiated at 60°C, ramped to 300°C at a rate of 10°C min⁻¹, and held at 300°C for 6 minutes, yielding a total analysis time of 29 minutes. Mass spectrometry detection was performed in electron ionization (EI) mode with a scan range of 45–900 m/z. The transfer line and ion source temperatures were both maintained at 230°C to facilitate efficient ionization and analyte transmission to the mass analyzer.

### Protein Extraction

Harvested bacterial pellets were suspended in 700 µL of Lysis buffer composed of 1% SDS, 50 mM Tris-HCl (pH 6.8), and 1 X working concentration of Halt™ Protease Inhibitor cocktail (Thermo Scientific, catalog no. 78430, Waltham, MA, USA). Cell lysis was sequentially performed by sonication and boiling. Using an appropriate microcentrifuge tube probe, pellet suspensions were sonicated at 20% power for 1 minute, with interval chilling on ice for 1 minute, this cycle is repeated five times. Post-sonication, tubes were placed in a heating block set at 90°C for 15 minutes. Tubes were centrifuged to separate cell debris, and lysate suspension was stored in a fresh protein Lo-Bind microcentrifuge tube. Crude protein lysate samples were stored at - 20°C. Crude protein lysates were quantified by absorbance spectrophotometry at 280nm wavelength. Crude protein lysates were treated using the Cytiva 2D Clean-Up Kit (Sigma-Aldrich code: GE80-6484-51). Protein mass (µg) was initially quantified from the estimated protein concentrations (µg/mL) generated by spectrophotometry. Protein volumes required for uniform protein masses (0.5 µg) were calculated prior to addition to 10 µL of SDS loading mix (5 µL Loading Buffer, 2 µL Reducing agent, 3 µL of Distilled H2O) and loading unto SDS-PAGE Lane, with a total load volume of 20 µL. Each protein mix were loaded onto a NuPAGE™ 4 to 12% Mini Protein gel for a short run of 5 minutes at 150V. SDS-PAGE gels were stained overnight using ReadyBlue© Coomassie Blue. Unresolved gel bands were then excised into 1 mm^3^ cube pieces and stored in Protein Lo-Bind© microcentrifuge tubes (PCR clean, catalog no. 022431081, Eppendorf AG, Hamburg, Germany).

### In-Gel Protein Tryptic Digestion

Gel bands were excised from the SDS-PAGE gel manually, post visualization by Coomassie blue stain. The gel bands were then de-stained and proteolytically digested with trypsin to generate peptides for mass spectrometry analysis using liquid chromatography LC–MS/MS as outlined by Shevchenko et al. <u>2006</u> (28).

### Liquid Chromatography Mass Spectrometry (LC-MS/MS) analysis

Peptide separation was achieved by reverse-phase HPLC with two mobile phase gradient system, using an C18 column (EASY-Spray PepMap RSLC; 50 cm × 75 μm ID, 2 μm; 40 °C; Thermo Fisher Scientific, Waltham, MA, USA). Solvent A (0.1% formic acid in water) and solvent B (0.1% formic acid in 80% acetonitrile) at a 300 nL/min flow rate (RSLCnano HPLC system; Thermo Fisher Scientific, Waltham, MA, USA), with a gradient program of 0–5 min at 6% B, increasing from 6% B to 50% B over the next 75 min. Mass spectrometry was performed using a Q Exactive HF hybrid quadrupole-Orbitrap (Thermo Fisher Scientific, Waltham, MA, USA). Two repeat injections were performed per sample analysed. Data-dependent acquisition was performed with 10 product ion scans (centroid: resolution, 30,000; automatic gain control, 1 × 105 maximum injection time, 60 ms; isolation: normalized collision energy, 27; intensity threshold, 1 × 105) per full mass spectrometry scan (profile: resolution, 120,000; automatic gain control, 1 × 106; maximum injection time, 60 ms; scan range, 375–1500 m/z).

### Metaproteomic Bioinformatics Workflow

Thermo (.raw) mass spectrum files for each tryptic peptide samples were converted into a mascot generic format (.mgf) via msconvert software. Each metapeptide databases for consortia BPS4 and BPS8 was concatenated with the common Repository of Adventitious Proteins (cRAP) contaminants library database (https://www.thegpm.org/crap/) prior to peptide spectrum matching. SearchGUI was used to match metapeptide databases (BPS4 and BPS8) against their respective mass spectrum chromatograms to identify distinct peptide peaks under the following parameters (enzyme: trypsin, max cleavage: 2, precursor ion tolerance: 10 ppm, fragment tolerance: 0.05 Da, fixed modifications: carbamidomeythylation of C, variable modifications: oxidation of M, validation at 1% false discovery rate [FDR]). PeptideShaker was used to generate peptide spectral matches (PSM), peptide, and protein reports, which were filtered to exclude identified contaminants from the cRAP database. Peptide sequences extracted from PSM reports were inputted into the Unipept (applications: pept2lca, peptinfo) to generate taxonomic, gene ontology, and enzyme commission datasets. Total PSM counts and PSM counts per identified genus ranks were generated via SQL between PSM reports and their respective taxonomic data output from Unipept (application: pept2lca). For Label-Free Quantification, an open source LFQ software FlashLFQ (Millikin et al., 2021) was used to extract the peak intensity from mass spectrum files. LFQ intensity data, in combination with extract Gene Ontology (GO) data and Enzyme Commission (EC) data from Unipept, were inputted into the metaQuantome an integrated metaproteomics software pipeline in GalaxyP, was used to quantify significant protein fold changes of annotated gene ontology and enzymes. All raw mass spectra have been submitted to the MassIVE repository (https://doi.org/doi:10.25345/C5JQ0T807).

## Results

### Characterization of Waste Oils

GC-MS analysis of WOs revealed distinct peaks that enabled the characterization of the complex petrochemical chemical mixture. Visualization of chromatogram (**Supplementary Figure 4**) shows 18 distinct peaks labeled for identification via matching against the NIST reference library. Each distinct peak was matched against its reference chemical chromatogram that scored with both high match and reverse Match following NIST’s general guidelines for match factors (**Table 1**). Of the identified and matched peaks, the chemicals that presented the highest of intensities were 10-Octadecenoic acid methyl ester (Peak #17), Benzene 1,2,4-trimethyl (Peak #3), Tributyle phosphate (Peak #11), and Hexadecanoic acid methyl ester (Peak #15). Overall GC-MS chromatogram showed a wide range of components starting from short-chain low molecular weight hydrocarbons such as Xylene (C-8) and Benzene (C-9), till long-chain high molecular weight hydrocarbons namely Eicosane (C-20) and Heptacosane (C-27). When comparing to annotated peaks with highest intensities visualized, intermediate-chain length hydrocarbons (C12-19) represented far lower peak intensities within WOs. Determination of PAH content (acetone/hexane extraction followed by GC-MS) identified that WOs contained about 313 mg/kg of PAHs in total, which is comprised of 16 different PAH components, which are listed by the United States Environmental Protection Agency (USEPA) as priority PAH pollutants (**Table 2**). Naphthalene, Phenanthrene, and Pyrene represent the most concentrated compounds within sampled WOs at masses of 84.5 mg/kg, 51.3 mg/kg, and 33.6 mg/kg, respectively.

**Table 1.** List of matched hydrocarbons from annotated peak scans in WOs sample against the NIST library via GC-MS.

| Peak # | Chemical Name | Retention Time<br>(minutes) | Match Score | R. Match Score |
| --- | --- | --- | --- | --- |
| 1 | P-Xylene | 2.4-2.5 | 742 | 900 |
| 2 | Benzene, 1-ethyl-3-methyl | 3.1 | 809 | 911 |
| 3 | Benzene, 1,2,4-trimethyl | 3.5 | 788 | 845 |
| 4 | Benzene, 1-ethyl-3-methyl | 3.8 | 757 | 891 |
| 5 | Hexadecane | 4.7 | 848 | 898 |
| 6 | Heptacosane | 6.1 | 727 | 764 |
| 7 | Tetradecane,<br>2,6,10-trimethyl | 7.4 | 816 | 843 |
| 8 | Tetradecane | 8.7-8.8 | 796 | 887 |
| 9 | Nonadecane | 10.0 | 804 | 921 |
| 10 | Hexadecane | 11.2 | 807 | 892 |
| 11 | Tributyl phosphate | 11.7-11.8 | 894 | 929 |
| 12 | Heptadecane | 12.3-12.4 | 855 | 936 |
| 13 | Phosphoric acid,<br>dibutyl phenyl<br>ester | 14.2 | 793 | 859 |
| 14 | Heptacosane | 14.4-14.5 | 820 | 859 |
| 15 | Hexadecanoic<br>acid, methyl ester | 14.7 | 849 | 882 |
| 16 | Eicosane | 15.4-15.5 | 823 | 904 |
| 17 | 10-Octadecenoic<br>acid, methyl ester | 16.4 | 845 | 852 |
| 18 | Heptacosane | 17.3 | 802 | 863 |

**Table 2.**
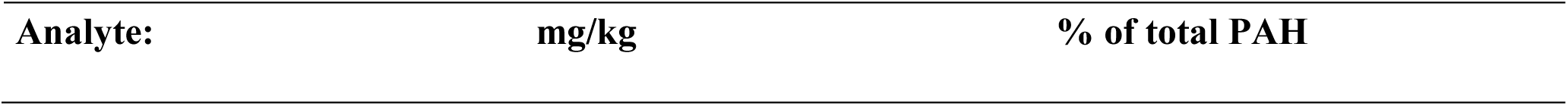
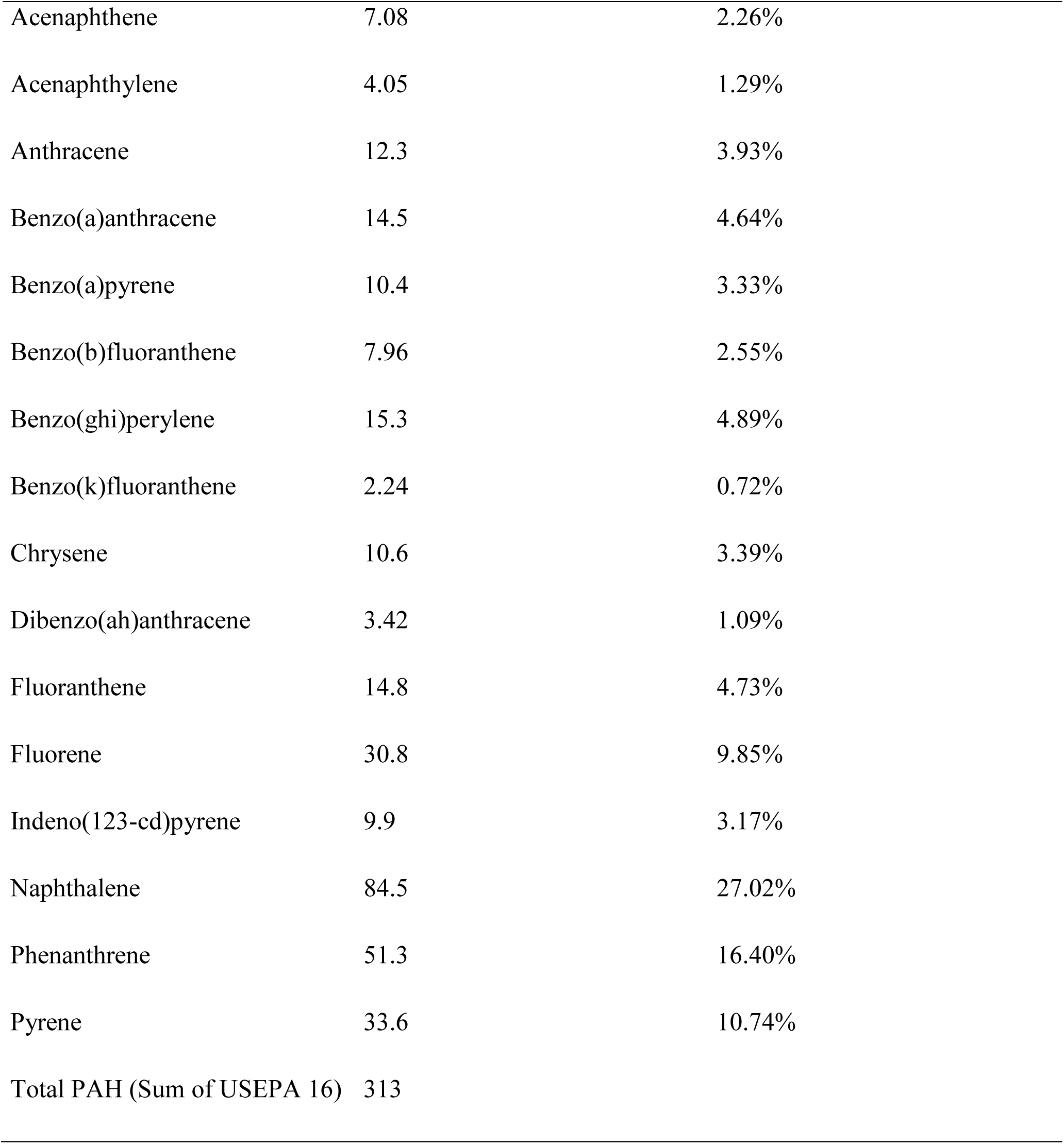
List of PAH content in WOs where mass of aromatics is quantified (mg) per kilogram of WOs.

| Analyte: | mg/kg | % of total PAH |
| --- | --- | --- |
| Acenaphthene | 7.08 | 2.26% |
| Acenaphthylene | 4.05 | 1.29% |
| Anthracene | 12.3 | 3.93% |
| Benzo(a)anthracene | 14.5 | 4.64% |
| Benzo(a)pyrene | 10.4 | 3.33% |
| Benzo(b)fluoranthene | 7.96 | 2.55% |
| Benzo(ghi)perylene | 15.3 | 4.89% |
| Benzo(k)fluoranthene | 2.24 | 0.72% |
| Chrysene | 10.6 | 3.39% |
| Dibenzo(ah)anthracene | 3.42 | 1.09% |
| Fluoranthene | 14.8 | 4.73% |
| Fluorene | 30.8 | 9.85% |
| Indeno(123-cd)pyrene | 9.9 | 3.17% |
| Naphthalene | 84.5 | 27.02% |
| Phenanthrene | 51.3 | 16.40% |
| Pyrene | 33.6 | 10.74% |
| Total PAH (Sum of USEPA 16) | 313 |  |

### Metagenomics: Taxonomy and Functional Prediction

Genomic DNA sequences of BPS4 resulted in a total of 185 million reads, of which 83% were taxonomically classified. In addition, 54.03% of total reads were classified at a species level, where normalized sequence reads showed an estimated abundance of 59% for *Stenotrophomonas maltophilia,* 21% for *Pseudomonas fluorescens,* 8% for *Delftia acidovorans,* and 3% for *Pseudomonas putida,* with the remaining classified species representing about 1% or less in abundance (**Figure 1**). Metagenomic sequence data functionally predicted through the Prodigal algorithm annotated with protein functionality, were filtered for enzymes involved with aliphatic and aromatics hydrocarbon compounds found in our WOs. Enzymatic pathways were reconstructed through literature and archived databases such as the International Union of Biochemistry and Molecular Biology (IUBMB) Enzyme Nomenclature, ExplorEnz, and the Kyoto Encyclopedia of Genes and Genomes (KEGG) (**Figure 5**). Out of the total of functionally annotated proteins 257 entries were associated with enzymatic pathways of hydrocarbon biodegradation. This included ring hydroxylating enzymes for aromatic hydrocarbons, such as toluene dioxygenases, naphthalene dioxygenases, and terminal oxidation enzymes that initiates the catabolism of aliphatic hydrocarbons, like alkanes monooxygenases. Catabolic reactions of aliphatics and aromatics both lead to the production of intermediates, specifically alkanes are targeted through terminal oxidation via alkane monooxygenases, and subsequently end with the production of fatty acyl-CoA that enters β-oxidation; while aromatics are initially targeted by ring hydroxylating oxygenases that consequently produce intermediate metabolites like catechols and phthalates, which are metabolized further into citric acid and succinyl-CoA that enter the TCA cycle. Enzymes located downstream of catechol and phthalate intermediates were found to be more widespread between metagenomic bins in comparison to monooxygenase and dioxygenases that initiates the cascade of petroleum hydrocarbons catabolism.

**Fig. 1.**
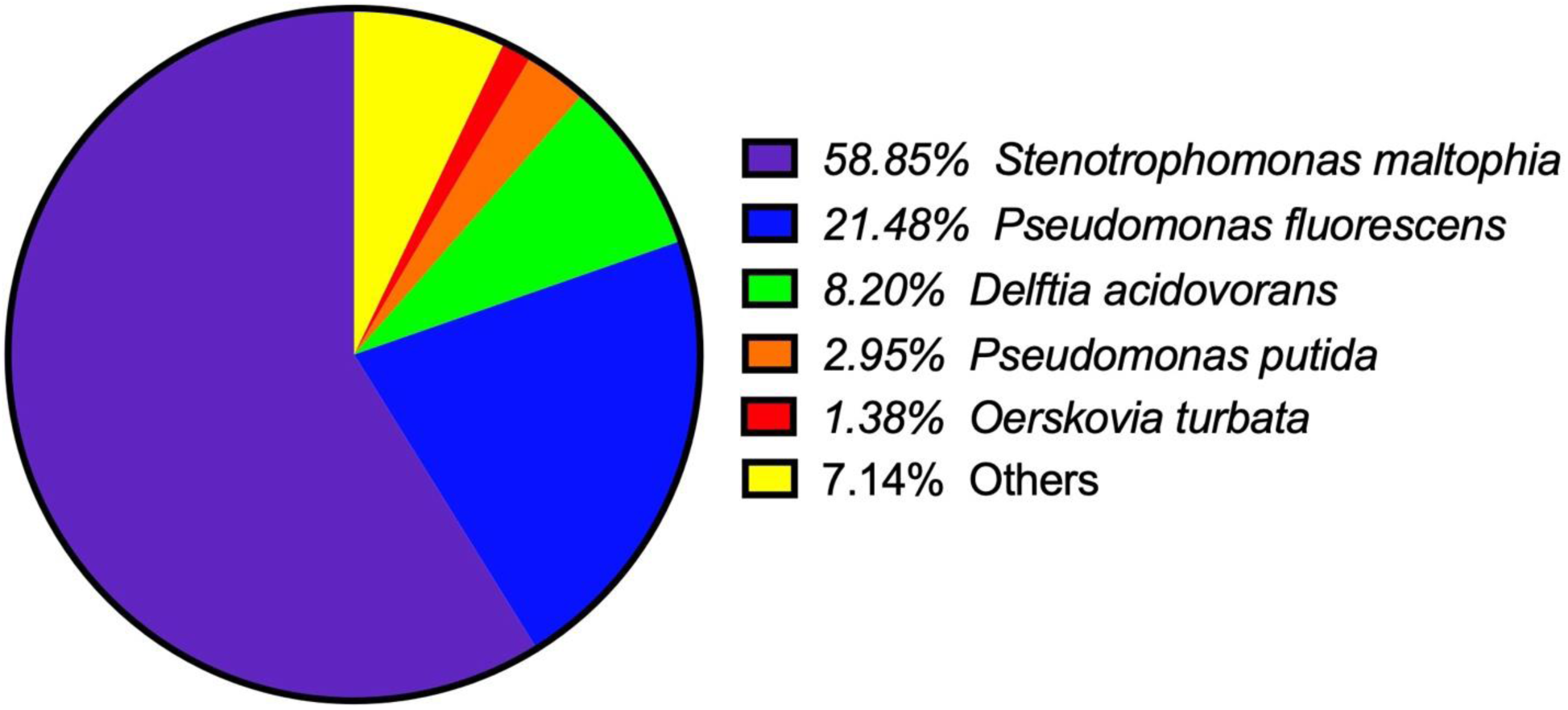
Taxonomic Abundance at the species level of normalized metagenomic 16S rRNA sequence reads for consortia BPS4. Species with less than 1% relative abundance were grouped into “Others” category.

### Metaproteomic taxonomy profile

Peptide spectral matching using Peptideshaker, and taxonomic annotation via Unipept, resulted with a total number of taxonomically annotated PSMs at an average of 26402 ± 41, 20609 ± 1940, 19147 ± 72, 19436 ± 932, and 10027 ± 270 between technical triplicate samples of A41, A42, O41, O42, and O43, respectively. Relative abundance of taxonomic annotations showed genus *Pseudomonas sp.* represented highly between 47%-52% of the bacterial community BPS4 when grown in acetate supplemented MSM media (A41, and A42), yet in oil supplemented media (O41, O42, and O43) it was less represented, between 32-36% of the community structure. In contrast, the genera *Delftia sp.* and *Achromobacter sp.* were less abundant in acetate MSM medium, comprising 14%-16% and 16%-20% of the community structure, respectively, while in WO supplemented medium they increase to 20% - 24% (*Delftia sp.*) and 26% - 27% (*Achromobacter sp.*). Less abundant genera were represented more uniformly across both acetate and oil cultures, which includes *Stenotrophomonas* between 5% and 7%, while the remaining genuses *Brucella*, *Microbacterium*, *Oerskovia*, and *Pseudochrobactrum* were represented between 2% to 3% (**Figure 2**). Pearson correlation analysis showed that certain members of consortia BPS4, such as *Pseudomonas,* had significant positive correlation (|r|>0.8, *p-*value*<*0.0001) with multiple members like *Pseudochrobactrum, Stenotrophomonas,* and *Microbacterium*, meanwhile *Achromobacter* and *Delftia* are exclusively correlated between each other within consortia BPS4. Though a noticeable variation in the overall taxonomic structure of genera is visualized in oil cultures (**Figure 2**), correlation coefficient intervals did not show negative correlations between taxonomic members, which was further observed in a principal component analysis (PCA) that showed a linear relationship with less than sufficient PC2 variation to be significant.

**Fig. 2.**
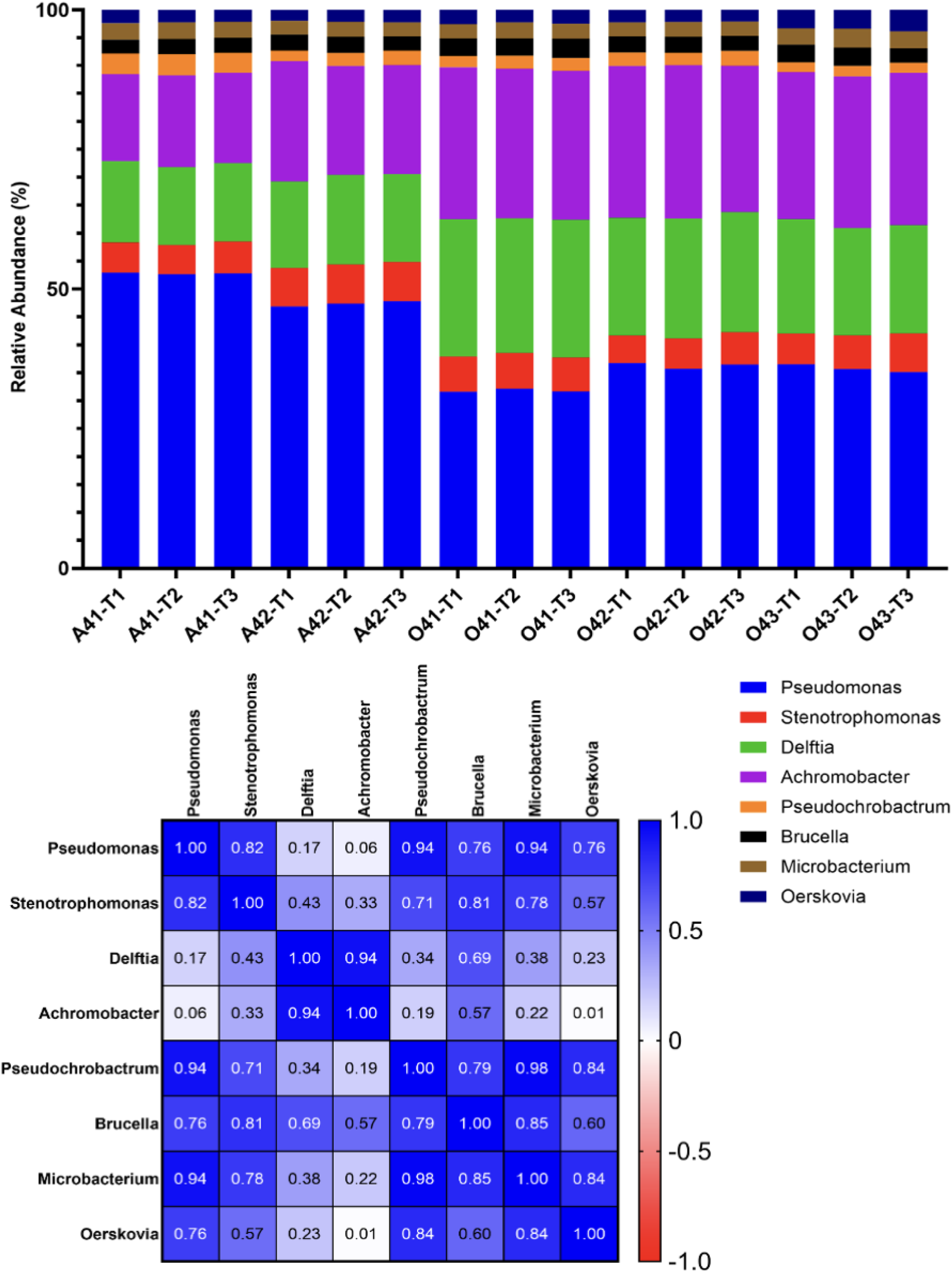
Relative abundance of taxonomically annotated peptides via PSM counts per identified genus (Top). Pearson correlation matrix between identified genera within consortia BPS4 (Bottom). Acronyms: A = acetate cultures; O = waste oil cultures; 41,42,43 = biological replicates of consortia BPS4; T1,T2,T3 = technical replicates.

### Gene Ontology enrichment

Each annotated GO ontology terms is determined based on the presence of peptide intensities that belong to the same protein group. GO annotated peptides were retained if peptide intensity was quantified in at least two of the triplicate technical replicates MS spectral data. From the generated protein groups, a total number of 6143 GO terms were annotated, of which 2971, 740, and 2431 are categorized under biological process, cellular component, and molecular function GO namespaces, respectively. PCA clustering was used to determine the degree of separation of GO annotated peptides, where the ratio of the average squared distance between all pairs of cluster centers over the sum of squared distances between each point and its respective cluster center, resulted with a moderate cluster separation of 77.3% (**Figure 3**). Differentially enriched GO terms between acetate and WO cultures were further explored by plotting peptide level intensity fold changes and selecting for significant terms (±1 Log_2_ peptide fold change, and ≤ 0.05 *P* value [-Log(FDR)] (**Figure 3**). In total, 227 biological process and 236 molecular functions terms plotted passed the fold-change and *p-*value cut-off. Within the top 30 significantly enriched GO terms for biological processes, the data showed several terms associated with hydrocarbon biodegradation, such as naphthalene catabolic process (GO:1901170), alkene metabolic process (GO:0043449), and cellular detoxification (GO:1990748). Under molecular functions, oxidoreductase activity terms were enriched in WO cultures that are associated with hydrocarbon biodegradation pathways, such as benzene 1,2-dioxygenase activity (GO:0018619), benzaldehyde dehydrogenase (NAD+) activity (GO:0018479), alcohol dehydrogenase (cytochrome c) activity, and other broader oxidoreductase activity terms. In both biological process and molecular functional GO annotated peptide intensities, nitric oxide biosynthesis (GO:0006809) and nitrate reductase activity (GO:0009703) were the significantly enriched in WO conditions, indicating high nitrogen cycling and/or regulatory activity within the community, including terms the cellular response to nitrogen starvation (GO:GO:0006995) and cellular response to nitrate (GO:0071249).

**Fig. 3.**
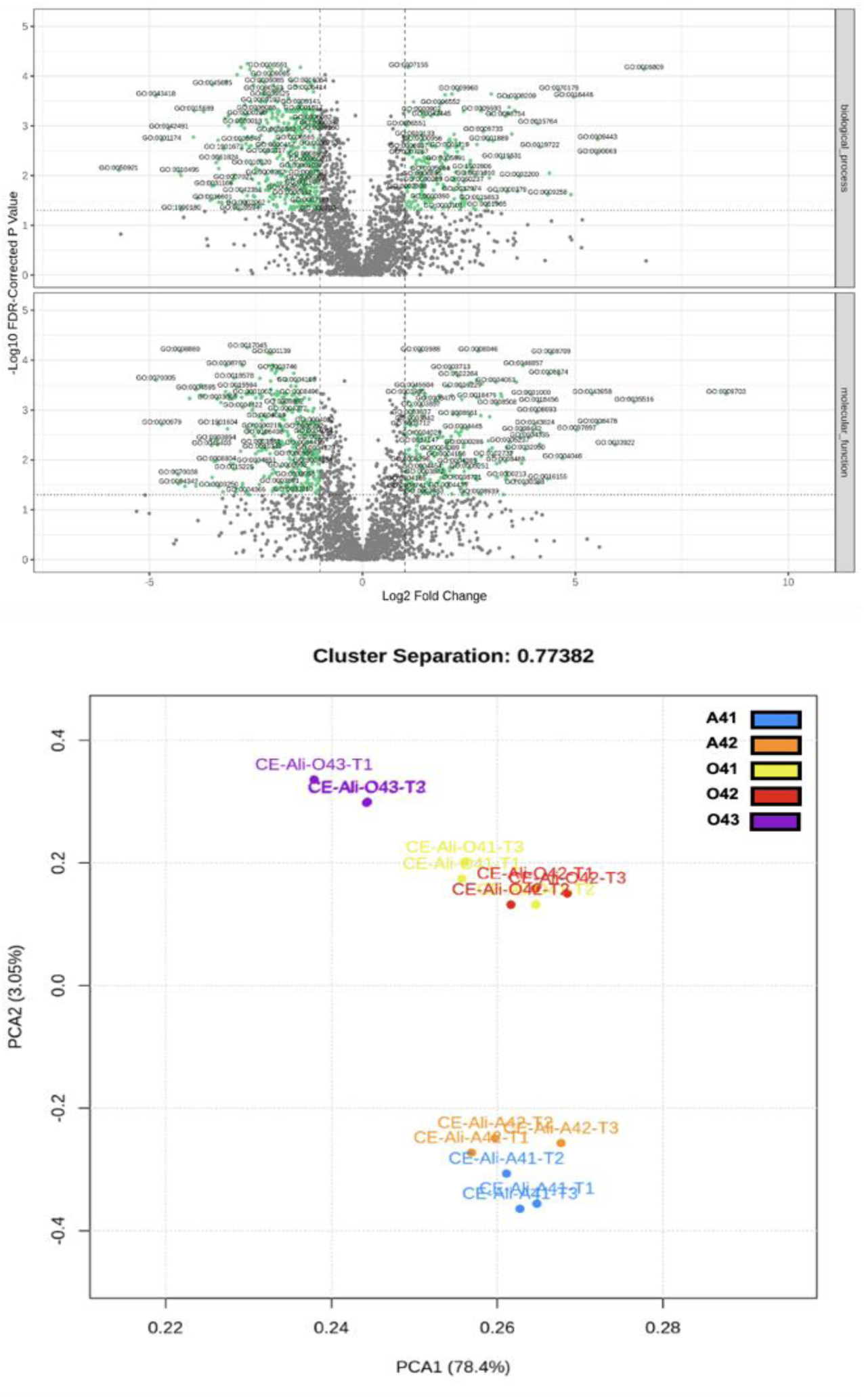
Volcano plot showing peptide-level fold change of significant GO annotations (cut-off = ±1 Log2 fold change, and p-value ≤ 0.05) (Top). PCA analysis of significant GO annotations per sample (Bottom). Acronyms: A = acetate cultures; O = waste oil cultures; 41, 42, 43 = biological replicates of consortia BPS4; T1, T2, T3 = technical replicates.

### Differential Enzyme Commission enrichment

A total number 1399 EC numbers were annotated for enriched peptides, where 392, 595, and 410 annotations in biological replicates of WO cultures (O41, O42, and O43, respectively) were validated with significant fold change (cut-off = ±1 Log_2_ fold change) and low FDR (*p* ≤ 0.05). In addition, PCA analysis showed EC annotated peptides between acetate and WO cultures with a moderate cluster separation of 77.7%. Mean peptide intensities show distinct clusters of EC annotations enriched between acetate and WO cultures, the majority of which 144 are represented under oxidoreductases, 147 transferases, and 136 hydrolases. EC annotations were filtered and selected for those present on the predicted hydrocarbon biodegradation pathways within consortia BPS4. EC annotations were grouped into aliphatic or aromatic hydrocarbon biodegradation pathway, with a cluster analysis of differentiated abundance of annotated peptide intensities between acetate and WO cultures (**Figure 4**). In total, EC annotated peptide fold changes showed 22 significant (*p*-value ≤ 0.05) differentially enriched enzymes in WO cultures associated with aliphatic biodegradation pathway, while aromatic pathway associated ECs resulted with 10 annotations that were significant. ECs under aliphatic hydrocarbon degradation showed enrichment of enzymes in WO cultures, such as two types of pyrroloquinoline quinone (PQQ)-dependent alcohol dehydrogenases, azurin (EC 1.1.9.1) and cytochrome (EC 1.1.2.8) by 2 fold, short chain acyl-CoA dehydrogenase (EC 1.3.8.1) by 4 folds, while other precursor enzymes that target longer chain acyl-CoA (EC 1.3.8.7 and EC 1.3.8.8) being less prevalent in WO cultures by 1.5 to 2 folds. Under the aromatic hydrocarbon biodegradation pathway classification, WO cultures were enriched with aryl-alcohol dehydrogenase (between 2.5 to 4 folds) and benzaldehyde dehydrogenase (between 3.5 to 4.0) fold.

**Fig. 4.**
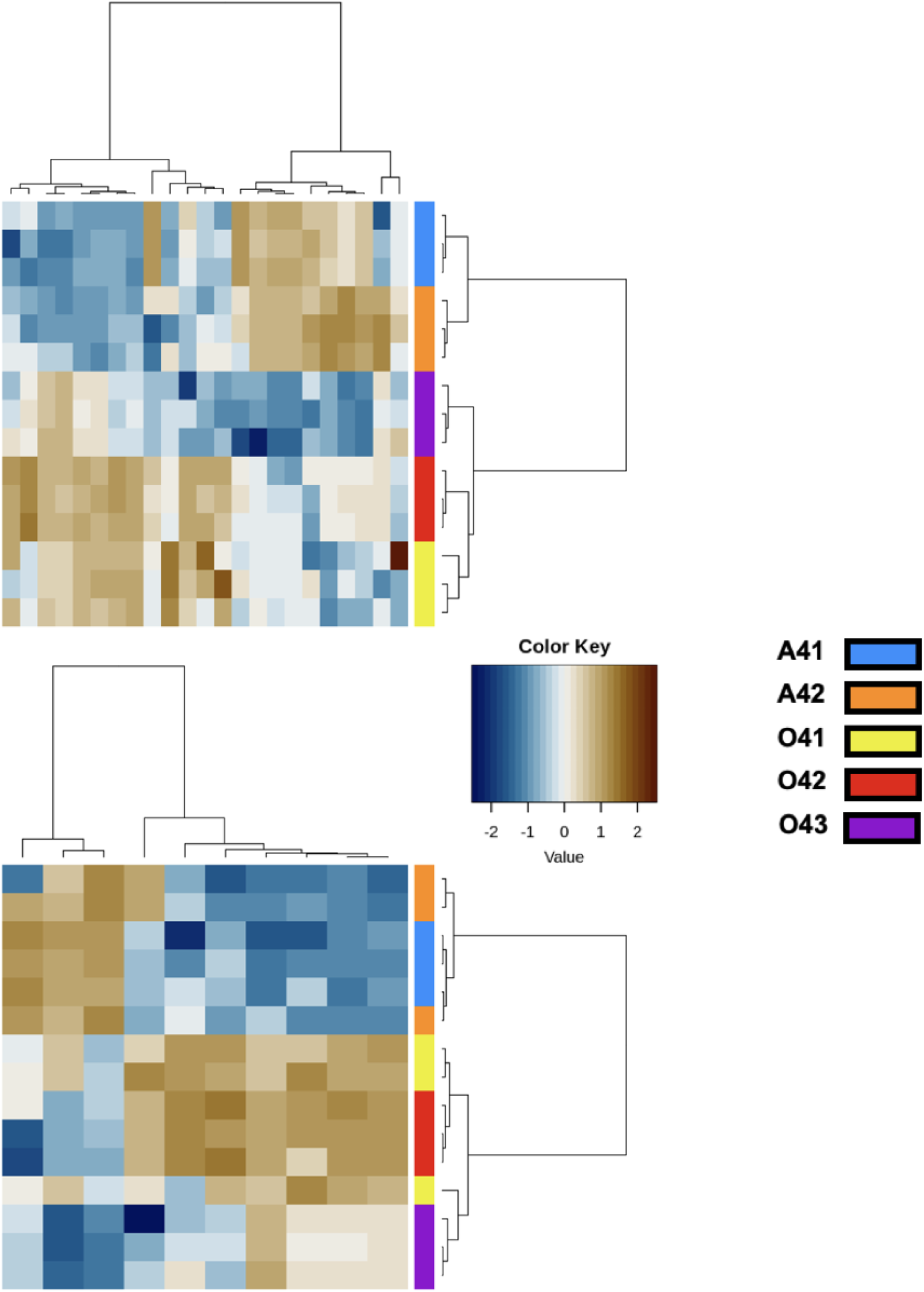
Differential enzyme activity profiles in hydrocarbon biodegradation pathways of consortium BPS4. Heat map showing differential abundance of significant enzyme commission (EC) annotations across BPS4 biological replicates in WO (O41, O42, O43) versus acetate control (A41, A42) cultures. Enzymes are grouped into aliphatic (upper panel) and aromatic (lower panel) hydrocarbon degradation pathways. Color intensity represents log₂ fold-change values, with 22 enzymes significantly enriched in aliphatic pathways and 10 in aromatic pathways (p ≤ 0.05).

### Label-Free Quantification & Hypothetical Protein Analysis

LFQ protein fold change analysis revealed 514 protein groups, of which 172 unannotated hypothetical proteins enriched in WO cultures (fold-change ≥ 1.5, at an FDR of ≤ 0.002). Each hypothetical protein group identified was cross referenced with the BPS4 metapeptide database to extract amino acid sequences queried into the NCBI BLAST-P tool for further taxonomic & functional annotation. Through the NCBI non-redundant (NR) protein database, significant protein (i.e. data entries at fold-change ≥ 1.5, and an FDR of ≤ 0.002) resulted with taxonomic annotations at a species level that affirmed genera previously annotated via the Unipept tool. A subset of highly abundant significant proteins associated with stress response show majority of entries under the genera *Achromobacter*, *Delftia*, and *Pseudomonas*, in contrast to other less abundant members such as *Oerskovia*, *Stenotrophomonas*, and *Brucella.* BLAST-P functional annotations found homologs of a histidine kinase sensor gene *RcsC* shared between *Achromobacter*, *Delftia*, *Pseudomonas,* and *Stenotrophomonas,* which were annotated as regulatory response proteins via the NCBI NR database. Annotated hypothetical proteins show homologs of ABC transporter substrate-binding proteins between *Delftia*, and *Achromobacter,* while some entries returned with no known physiological function, notably *Pseudomonas putida* (WP_115272915.1), *Achromobacter xylosoxidans* (CUJ85469.1), and *Oerskovia sp.* (WP_336708573.1).

### Waste oil biodegradation: Post Batch-fed Adaptation

GC-MS analysis of extractable residual WOs from adapted cultures (ROS-AD) were compared to unadapted (ROS-UN) cultures to assess performance of the batch-fed protocol in enhancing biodegradation capabilities of consortia BPS4. Untreated WO (ROS-1g) exhibited a chromatogram featuring 18 resolved peaks corresponding to a wide distribution of intermediary to long range n-alkanes (e.g., tetradecane, pentadecane, hexadecane, heptadecane, octadecane, eicosane, etc.) and branched or functionalized hydrocarbons such as 2,6,10,14-tetramethylpentadecane and methyl esters (**Figure 6**; **Table 3**). Biologically treated samples (ROS-UN and ROS-AD) displayed marked depletion of resolved n-alkanes, evidenced by greatly diminished or absence of peaks across the lower and middle range retention times (**Figure 6**; **Table 3**). Compositional difference was further substantiated by compound identification where only recalcitrant components such as 2,6,10,14-tetramethylpentadecane, 2,6,10,14-tetramethylhexadecane, 9-methylnonadecane, and phosphate esters (dibutyl phenyl phosphate, tributyl phosphate) were consistently detected in ROS-UN and ROS-AD (**Table 3**). Based on the chromatographic and compound identification data provided, there are minimal observable differences between ROS-UN (unadapted bacterial consortia) and ROS-AD (adapted bacterial consortia) in terms of hydrocarbon degradation patterns. Both samples exhibit nearly identical compound retention patterns, with six compounds detectable (**Table 3**). The chromatograms show comparable unresolved complex mixture patterns with broad, elevated baselines and similarly diminished peak intensities relative to ROS-1g. Overall, bacterial consortia demonstrated nearly complete removal of n-alkanes (C14-C28) and fatty acid methyl esters that were present in the untreated ROS-1g sample.

**Table 3.** Residual WOs identified by GC-MS in untreated (ROS-1g) and biologically treated (ROS-UN and ROS-AD) WO samples. Presence (+) or absence (–) of each matched peak compounds are indicated for unadapted (ROS-UN) and adapted (ROS-AD) experimental conditions.

| Chemical Name (Retention Time<br>[minutes]) | ROS-1g | ROS-UN | ROS-AD |
| --- | --- | --- | --- |
| Tetradecane (9.7) | + | - | - |
| Pentadecane (10.9) | + | - | - |
| Hexadecane (12.1) | + | - | - |
| Tributyl Phosphate (12.9) | + | + | + |
| Heptadecane (13.1) | + | - | - |
| 2,6,10,14-Tetramethylpentadecane (13.2) | + | + | + |
| Octadecane (14.2) | + | - | - |
| 2,6,10,14-Tetramethylhexadecane (14.3) | + | + | + |
| Nonadecane (15.1) | + | - | - |
| 9-Methylnonadecane (15.3) | + | + | + |
| Dibutyl phenyl phosphate (15.4) | + | + | + |
| Hexadecanoic acid, Methy Ester (15.6) | + | - | - |
| Eicosane (16.1) | + | - | - |
| Heneicosane (17.1) | + | - | - |
| Oleic acid, Methyl Ester (17.2) | + | - | - |
| Docosane (17.8) | + | - | - |
| Octacosane (18.7) | + | - | - |

### Consortia Metaproteome: WO Batch-fed Adaptation

Notable differentiated data entries between WO cultures of BPS4 ROS-UN and ROS-AD revealed distinct functions that were not shown in previous metaproteome analysis of WO cultures. Under gene ontologies enrichment, exhibited increased abundance of Rho protein signal transduction (5.13-fold), dodecyl sulfate metabolism (2.88-fold), copper ion import (3.49-fold), and multiple cellular signaling processes. At the molecular function level, notable entries included phosphoenolpyruvate mutase activity (3.48-fold), linear primary-alkylsulfatase activity (2.88-fold), long-chain fatty acyl-CoA hydrolase activity (2.60-fold) and phosphatidylcholine binding (2.56-fold). Differential analysis of enzyme annotations increased fold change in ROS-AD included ferric-chelate reductase (EC 1.16.1.9) at a fold change of 4.57, followed by phosphoenolpyruvate mutase (EC 5.4.2.9) with a fold change of 3.48. Linear primary-alkylsulfatase (EC 3.1.6.21) exhibited substantial upregulation (3.21-fold), alongside succinyl-CoA--L-malate CoA-transferase (EC 2.8.3.22) showing a fold change of 2.12. The coproporphyrinogen III oxidase (EC 1.3.3.15) and protein disulfide-isomerase (EC 5.3.4.1) displayed moderate increased abundance with fold changes of 1.96 and 1.84, respectively. Label free quantification for protein group entries with increased abundance in ROS-AD resulted with a total of 769 proteins of high confidence (FDR ≤.0.002) (**Figure 7**), of which, 500 proteins were functionally annotated while 269 were hypothetical or putative in functions.

**Fig. 5.**
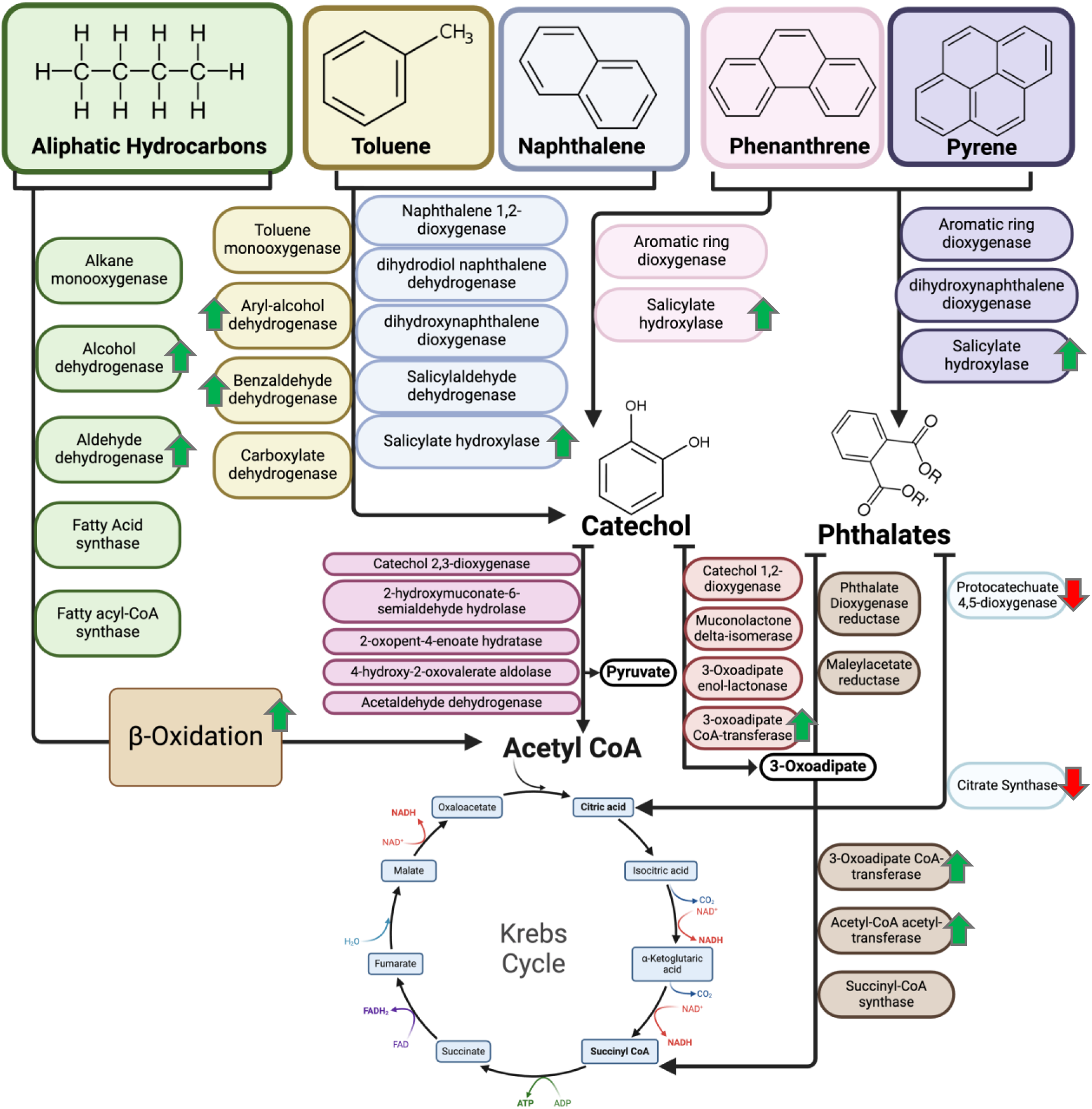
Illustration of predicted enzymatic pathways in consortia BPS4 involved with the catabolism of aliphatic, and aromatic hydrocarbons, which includes intermediate metabolites that are metabolized into the TCA cycle, based on literature, and archived enzyme commission databases. Green arrows annotated indicated increased differential abundance, while red arrows indicated reduced abundance of enzymes in WOs cultures compared to acetate cultures, as reflected from EC and LFQ analysis of metaproteomes.

**Fig. 6.**
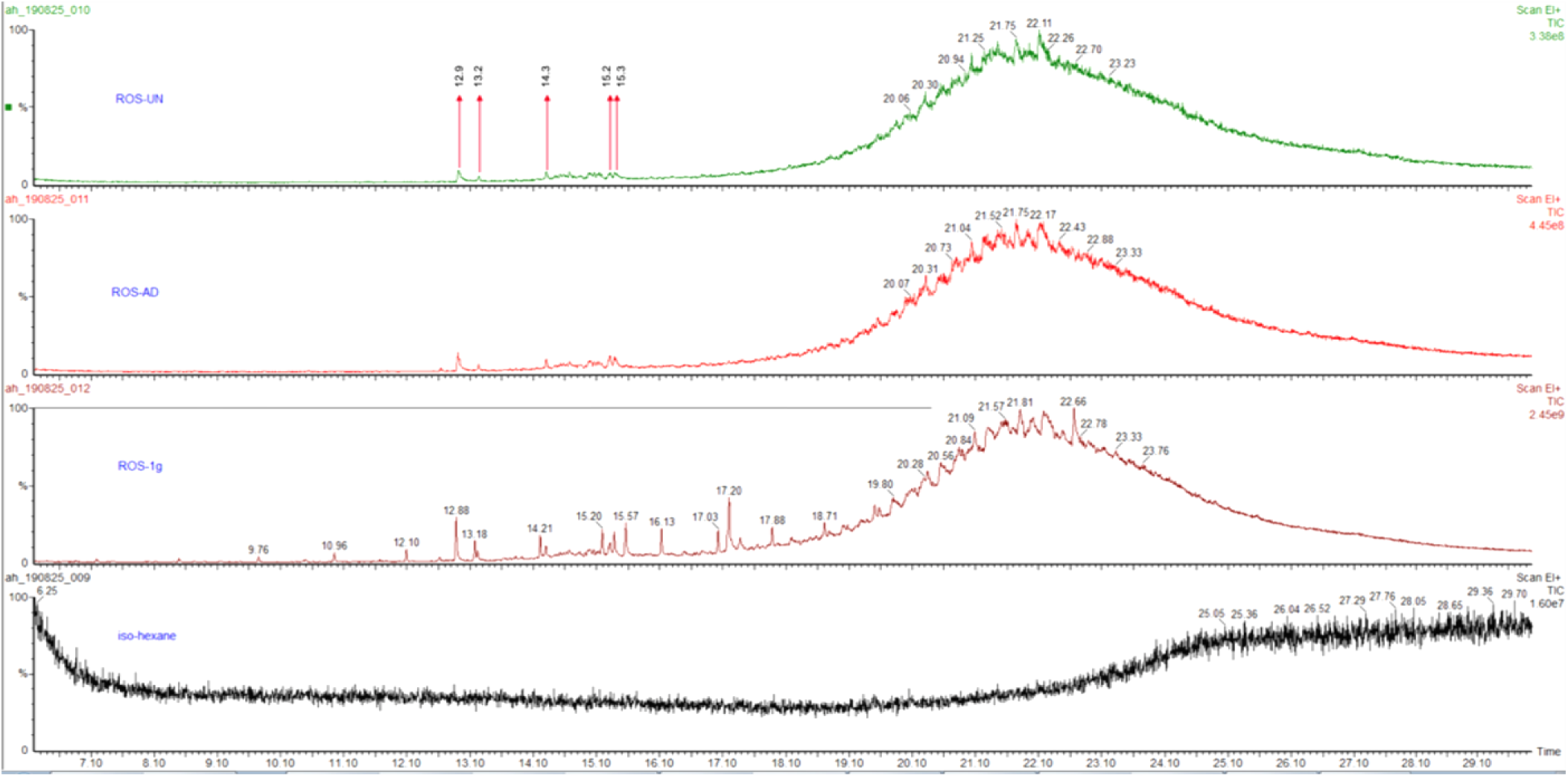
GC-MS total ion chromatograms of untreated WO (ROS-1g), WOs after biological treatment with unadapted (ROS-UN) and adapted (ROS-AD) bacterial consortia, and iso-hexane solvent blank. Recalcitrant peaks are labeled with retention times for representative compounds.

**Fig. 7.**
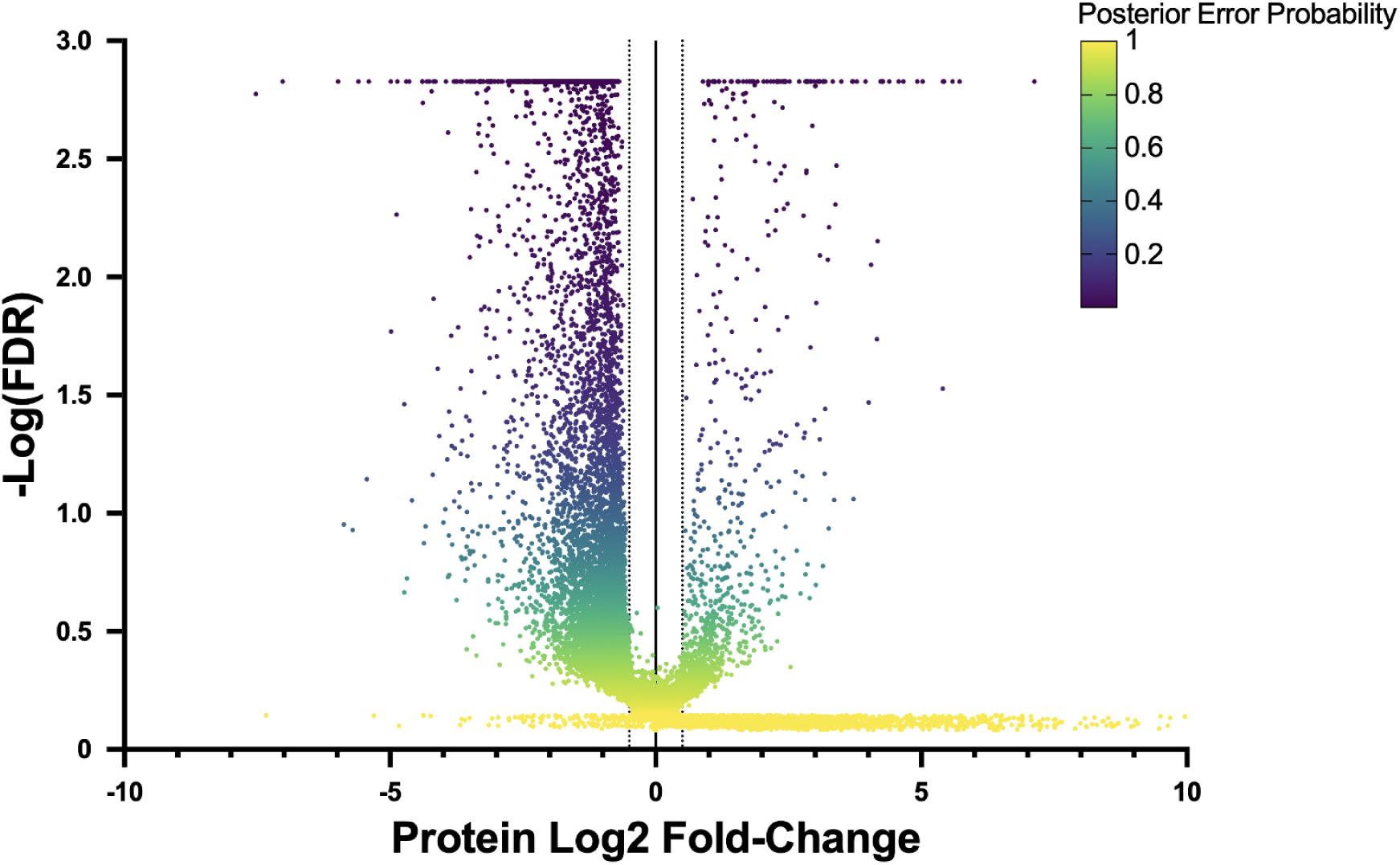
Volcano plot of LFQ protein quantifications showing fold-change (cut-off = ±1.5) of significant protein entries (FDR of ≤ 0.002, Posterior Error Probability ≤ 0.2) for cultures ROS-UN versus ROS-AD of bacterial consortia BPS4.

Of the functionally annotated and hypothetical protein group entries, 54 and 21 were presented with a protein Log_2_ fold change of ≥ 3. Significant findings included multiple hits associated with alkyl/arylsulfatases taxonomically annotated with *Achromobacter* and *Pseudomonas* with a fold change between 5.02 to 7.13. Additional metabolic processes identified were grouped into uridine kinase from *Oerskovia, a*lpha-ketoacid dehydrogenase subunit from *Achromobacter,* and a functionally uncharacterized protein from *Delftia* (5.72-fold). Other metabolic functions included 2,5-didehydrogluconate reductase, and an aldo/keto reductase from *Achromobacter* (4.23-fold), and a class I SAM-dependent methyltransferase from *Pseudomonas* (3.49-fold). Transport functions were also shown such as amino acid ABC transporter-binding proteins from *Achromobacter* (3.01-fold) and ATP-binding cassette domain-containing proteins from *Streptomyces* (3-fold). Interestingly, a FliI/YscN family ATPases was annotated and taxonomically assigned for *Oerskovia* and *Microbacterium* with a fold change of 4.57, despite representing minor members of the consortium’s relative abundance of taxons. Two hypothetical protein entries (database accessions: GEEJPKPN_77088 and GEEJPKPN_02100) did not present significant hits against the ClusterNR, Swissprot, and Refseq databases.

## Discussion

### Enrichment strategy for HDB consortia

One aspect that makes a top-down approach more appealing for enrichment is its less strenuous labor, especially when compared to conventional process of enriching and isolating individual microbial colonies, and screening for members of interest. Regardless, many studies have successfully constructed bacterial, fungal and micro-algae consortia with 2 or more distinct species, including more diverse co-kingdom communities for the bioremediation of petroleum hydrocarbons (29–31). With more complex compositions of microbial consortia, implementing conventional bottom-up approaches of enrichment can become cumbersome, including the challenge of stabilization and control of such complex consortiums. Christensen et al. (2002) found two species, *Pseudomonas putida* strain R1 and *Acinetobacter* strain C6, where the latter member was dominant for the metabolism of benzyl alcohols as a sole carbon source (32). However, microniches within the biofilm space were found to circumvent competition into commensalism, where upper layers of biofilms dominated by *Acinetobacter* strain C6 metabolizing benzyl alcohols lead to excess benzoate diffusing into outer vicinities into the *Pseudomonas putida* R1 occupied space allowing it to cooperatively metabolize rather than compete for the precursor benzyl alcohols (32). Despite the challenges, meta-analysis studies suggest application of microbial consortia are more favorable to single species inoculation (33). Although bottom-up enrichment is not without merit, as utilizing top-down enrichment to inform bottom-up isolation has proven crucial for understanding and constructing syntrophic consortia (11).

Soil enrichment cultures from waste oil contaminated sites BPS4 and BPS8 provided two consortia with recognizable variation of abundance of bacterial genera. Enriched consortia predominantly belong to the phylum Pseudomonadota (also known as Proteobacteria), which conforms with many prokaryotic hydrocarbon biodegradation studies (34). Metagenomic profiling showed *Stenotrophomonas maltophilia* to be the dominate species in relative abundance within both enriched consortia. Less abundant members found within enriched consortiums were *Pseudomonas fluorescens, Delftia acidovorans,* and *Pseudomonas putida* (consortia BPS4); *Stenotrophomonas terrre*, *Achromobacter piechaudii*, and *Stenotrophomonas humi* (consortia BPS8). Metagenomic results showed a more diverse abundance of species in consortia BPS4, where *P. fluorescens, D. acidovorans,* and *P. putida* cover a relative abundance total of 33%, notable for their petroleum hydrocarbon biodegradation. Our enrichment technique resulted with a community of ubiquitous bacteria found in a variety of environments and found to tolerate and degrade xenobiotics. *S. maltophilia* is characterized as a tolerant to xenobiotics, especially to heavy metals (copper and zinc) in tandem to degradation of C10-C34 petroleum hydrocarbons (35). Recent study recognized *S. maltophilia* as a biosurfactant producer capable of improving bioavailability of petroleum oil biodegradation for other members within a microbiome (36). In hydrocarbon biodegradation, the *Pseudomonad* group of bacteria are readily found in soil and water environments. Notably, *P. putida* is renowned for its metabolic versatility, non-pathogenicity, and capability of tolerating physiochemical stresses along with degradation of toxic xenobiotics (37). *D. acidovorans* (previously known as *Comamonas acidovorans*) (38) was previously enriched using phenanthrene as a sole carbon source, and their mechanism of biodegradation pathway was unraveled via whole genome sequencing (39). Despite metagenomic sequencing data aligning with previous reports identifying hydrocarbon-degrading bacteria, the metaproteomic taxonomic analysis raises further questions regarding the taxonomic composition of microbial communities and their restructuring under hydrocarbon exposure at the protein level.

### Metaproteomics: filling community-function landscapes

Bottom-up proteomic analysis of consortia grown in acetate containing media versus hydrocarbon containing media revealed a marked difference in structure compared to initial metagenomic data. Metaproteomics data showed *Pseudomonas* as the dominant genus in the community, followed by *Achromobacter* and *Delftia*. The genera *Delftia.* (*Comamonadaceae* family) and *Achromobacter* (*Alcaligenaceae* family), belonging to the β-proteobacteria lineage, within consortia BPS4 show a relative increase in abundance under WO growth as a sole carbon source. More importantly, *Stenotrophomonas* was shown to cover significantly less relative abundance. Early in this study, taxonomic analysis assigned a majority of sequence reads under *Stenotrophomonas maltophia,* however metaproteomic data revealed a reclassification from the family *Xanthomonadaceae* to *Lysobacteracea*e, initially suggesting a discrepancy in our sequence data and database library generated. However, a recent study suggested 16S sequencing has low discriminately for related *Stenotrophomonas* species, where 1555 genomes annotated as the clinically notable pathogen *S. maltophilia*, 45.34% (705 strains) did not actually belong to this species (40). In combination with metaproteomic insights, our data suggests a lesser role for Stenotrophomonas within consortia BPS4, while *Delftia, Achromobacter,* and *Pseudomonas* exhibit more weight in the structure of consortia BPS4. Nonetheless, we attempt to transform our microbial consortia analysis from a static inventory of taxa into a dynamic, topographic map of metabolic activity through metaproteomics. A major challenge in community function landscape studies is functional redundancy, meaning distinct communities performing identical functions, a phenomenon that is theorized as an evolutionary protectant against functional extinction. For example, microplastic degrading biofilm studies showing distinct bacterial assemblages on polyethylene versus polystyrene may exhibit decoupled genomic and proteomic profiles; while their taxonomy differs, their expression of central carbon metabolism and stress response proteins remains identical (41).

Metaproteomic profiles often reveal that diverse communities converge on a shared metabolic expression profile by measuring the protein landscape independent of the taxonomic landscape. In our study, we focused on highlighting statistically significant gene otology, and enzyme commission peptide annotations through differential label free quantification. Several biological and molecular GO annotations with significant increase in the presence of oil as a sole carbon source play a role in the nitrogen availability cycle, including nitric oxide biosynthesis (6.6-fold), response to nitrogen starvation (3-fold), response to nitrate exposure (2-fold), and nitrate reductase activity (8.5-fold). Nitric oxide biosynthesis was presented as the top biological process GO for BPS4 in WO cultures, which may indicate high denitrification activity within the consortia. Despite the fact BPS4 was grown in aerobic conditions, in literature several key hydrocarbon degrading bacteria are dependent on nitric oxides as an electron acceptor, under anoxic and/or low-oxygen conditions, to degrade different types of PAHs and potentially playing a role in regulation of oxygenase activity (42). Nevertheless, differential abundance of functionally annotated peptides revealed a high nitrogen cycling in WO cultures, indicating favorable biodegradation conditions. This is supported by a study where heavy stable isotopes of nitrogen supplemented to freshly contaminated soils were found to be advantageous for membrane integrity of TPH degrading bacteria within a fungal-bacterial community, suggesting higher rate of nitrogen transformation rate can lead to more TPH conversion to assimilated carbon (43). Another study confirms the importance of the type of nitrogen source with biosurfactant production, where a strain of *Achromobacter sp.* exhibited significant biosurfactant production under (NH_4_)_2_SO_4_ supplementation as an inorganic nitrogen source (44).

Metaproteomic analysis revealed significant upregulation of pyridoxal 5’-phosphate (PLP) salvage and pyridoxal kinase activities in WO cultures, with a 5.5-fold increase in both. This coordinated elevation suggests that bacterial consortium members exhibit auxotrophy for vitamin B₆, which substantiated by the concurrent 3.6-fold enrichment of the GO annotation for folic acid-containing compound catabolic processes (GO:0009397). Pyridoxal kinase and the PLP salvage pathway function to maintain bioavailable PLP pools, which serve as essential cofactors in over 180 biochemical reactions fundamental to bacterial metabolism (45). In addition, a 4.6-fold elevation in aminoacylase activity was detected via integrated GO and EC fold-change analysis. This enzymatic enrichment facilitates the hydrolysis of N-acylated L-amino acid derivatives into their constituent L-amino acids and acyl moieties, enabling the consortium to process xenobiotic-derived amino acid conjugates (46). These metabolic adaptations were accompanied by substantial shifts in amino acid modification systems, including 4.6-fold increases in serine and threonine racemase activities and a 3.5-fold elevation in broad-spectrum D-amino acid biosynthesis. The functional significance of these modifications extends to multiple physiological roles: D-amino acid production modulates peptidoglycan biosynthesis through post-translational modifications, particularly via serine/threonine phosphorylation of cell wall-associated enzymes, while the incorporation of alternative D-amino acids into peptidoglycan stem peptides may represent a stress-response mechanism conferring resistance to β-lactam antibiotics (47, 48). These coordinated metabolic adjustments likely constitute adaptive responses to hydrocarbon-induced alterations in membrane fluidity, serving to maintain cellular membrane integrity. Overall, these findings indicate that the BPS4 consortium employs multifaceted metabolic strategies in response to WO exposure, including enhanced nitrogen cycling capacity, vitamin B₆ auxotrophic compensation mechanisms, and reinforced peptidoglycan remodeling to sustain membrane homeostasis under lipophilic stress conditions.

Functional prediction of hypothetical proteins is essential for elucidating metabolic networks, both direct and indirect, that participate in hydrocarbon biodegradation. It is important to acknowledge that BLAST-P sequence-based functional annotation, while rapid and cost-effective, does not account for tertiary protein structure and folding dynamics, which are factors essential for comprehensive functional characterization. Advanced structural prediction tools such as AlphaFold have emerged as powerful complementary approaches, particularly for hypothetical protein analysis (49, 50). A significant finding from our annotation analysis was the pronounced enrichment of ATP-binding cassette (ABC) transporter substrate-binding proteins (SBPs) in WO cultures. ABC transporter SBPs were highly enriched among members of *Delftia* sp. (GEEJPKPN_15010 [6.0-fold], GEEJPKPN_30493 [3.5-fold]) and *Achromobacter* sp. (GEEJPKPN_13188 [4.0-fold], GEEJPKPN_34286 [4.0-fold], GEEJPKPN_16905 [4.0-fold], GEEJPKPN_09141 [3.5-fold]). ABC transporter SBPs represent a critical, highly conserved component of prokaryotic transport systems, facilitating selective uptake of diverse substrates, including amino acids, peptides, sugars, metal ions, and specialized metabolites, particularly from nutrient limited environments (51). Prior investigation of *Rhodopseudomonas palustris* ABC transporter SBPs (n = 108 proteins) demonstrated substrate-specific differentiation via fluorescence thermal shift (FTS) assays, revealing functional interactions with transcriptional regulators of the MarR family (52). More recent studies have documented that ABC transporter SBPs exhibit substantial enrichment during hydrocarbon biodegradation, with particularly high abundance in biosurfactant synthesis pathways and unculturable microbial taxa recovered from petroleum-contaminated soils (53, 54). The coordinated enrichment of ABC transporter SBPs with the relative increase in *Delftia* sp. and *Achromobacter* sp. abundance suggests these proteins mediate critical substrate acquisition functions under hydrocarbon-induced selective pressure, despite functional annotations remaining incomplete for some hypothetical protein sequences.

### Preferential Biodegradation Pathways

Gene ontologies provide valuable frameworks for broad functional categorization, encompassing housekeeping metabolism, substrate transport, and stress response mechanisms. However, comprehensive characterization of defined metabolic pathways, particularly those mediating xenobiotic biodegradation, necessitates enzyme-level annotation to elucidate the biochemical pathway dynamics of microbial consortia under investigation. In this study, EC annotation analysis of peptides associated with aliphatic hydrocarbon degradation revealed significant enrichment in oil supplemented cultures, notably including two pyrroloquinoline quinone (PQQ)-dependent alcohol dehydrogenases: azurin (EC 1.1.9.1) and cytochrome c oxidoreductase (EC 1.1.2.8), each elevated approximately 2-fold. These oxidoreductases catalyze the sequential oxidation of primary alcohols generated via monooxygenase-mediated hydroxylation of alkanes to aldehydes, and subsequently to carboxylic acids, which feed into the β-oxidation cycle. This pathway enrichment is corroborated by the marked 4-fold elevation of short-chain acyl-CoA dehydrogenase (EC 1.3.8.1), which processes the C2–C6 acyl-CoA intermediates resulting from accelerated β-oxidation. In contrast, long-chain acyl-CoA dehydrogenases (EC 1.3.8.7 and EC 1.3.8.8) exhibited reduced representation, declining 1.5- to 2-fold in WO cultures relative to acetate controls, suggesting metabolic specialization toward lower-molecular-weight alkane degradation under the imposed selective pressure. Notably, PQQ-dependent alcohol dehydrogenases utilizing azurin or cytochrome c as electron acceptors have demonstrated utility in biotechnology applications, particularly in addressing metabolic limitations encountered in the development and industrial scaling of synthetic methylotrophic bacterial strains (55, 56).

Under the aromatic hydrocarbon biodegradation pathway, WO cultures exhibited coordinated enrichment of aryl-alcohol dehydrogenase (2.5 to 4.0-fold) and benzaldehyde dehydrogenase (3.5 to 4.0-fold), which catalyze the sequential oxidative conversion of toluene derivatives through benzoate and salicylate to catechol, key central intermediates in the aromatic degradation cascade (57). Phthalate metabolism represents an alternative pathway generating protocatechuate; however, protocatechuate 4,5-dioxygenase showed markedly reduced abundance in oil cultures, as did citrate synthase, indicating metabolic specialization toward catechol-dependent degradation pathways over protocatechuate-mediated routes. This enzymatic selectivity suggests preferential channeling of acetyl-CoA generated from catechol cleavage toward succinyl-CoA formation and the citric acid cycle, thereby optimizing carbon assimilation efficiency (**Figure 5**). The dominance of gram-negative bacteria within the BPS4 consortium, which preferentially utilize catechol intermediates over phthalate-derived protocatechuate for aromatic hydrocarbon biodegradation, further supports this metabolic partitioning (57). Collectively, the coordinated upregulation of initial oxidative enzymes coupled with selective enrichment of downstream pathway components reflects a metabolic strategy that balances substrate activation and carbon assimilation. The concurrent downregulation of alternative pathway branches indicates functional specialization that enables optimal resource allocation and metabolic efficiency under the imposed selective pressure of petroleum hydrocarbon degradation.

### Sustained Biodegradation Efficiency Following Adaptation

GC-MS analysis of metabolites from batch-fed-adapted consortia revealed minimal compositional differences between non-adapted control (ROS-UN) and adapted (ROS-AD) cultures, suggesting that the batch-fed enrichment process did not substantially enhance the degradative capacity of the bacterial consortia under the tested experimental conditions. Previous reports have documented significant improvements in degradation efficiency following microbial adaptation, for example, a study conducted by Arias-Sánchez et al. (2024) using an experimental design for artificial selection demonstrated improved microbial community functions for biodegradation of pollutants without substantial genetic evolution (58). *In-silico* modeling of microbial community dynamics similarly demonstrated that directed evolution can optimize consortium function through ecological mechanisms, such as metabolic syntrophy, niche portioning, or community successions rather than genetic modifications. Chang et al. (2021) showed that top-down synthesized consortia exhibit substantially greater resistance to invasion by external members compared to bottom-up synthesized consortia, suggesting that ecological assembly strategies may confer competitive advantages independent of genetic optimization (59). In our study, it is possible that the adaptation period would need to be extended, or culture conditions did not promote rapid evolution, like continuous cultivation may have achieved. In addition, it is possible that the consortia reached metabolic saturation under the imposed selective pressure. A notable secondary observation was the pronounced depletion of lower molecular weight hydrocarbons in adapted cultures relative to the baseline data presented initially, particularly the absence of benzene, toluene, ethylbenzene, and xylene (BTEX) compounds. BTEX compound depletion likely resulted from physical evaporation rather than microbial metabolism. High volatility and low molecular weight hydrocarbons are inherently susceptible to loss during prolonged storage with repeated vial transfers and under aerated batch culture conditions, where continuous aeration and agitation create a headspace environment conducive to vapor-phase loss. This technical limitation is well-documented, and specialized bioreactor designs incorporating volatility-mitigation strategies have been developed to address this constraint (60, 61).

Taxonomic analysis of ROS-UN and ROS-AD surprisingly showed stability with minimal taxonomic differences, with both maintaining nearly identical relative abundance patterns dominated by *Achromobacter* (48-52%) and *Pseudomonas* (25-30%). This suggests adaptation period was insufficient to drive taxonomic restructuring, bacterial community structure has already exhausted potential variance, or that the bacterial community’s functional redundancy has reached an optimal state for waste oil degradation. It is important to highlight consortia BPS4 revived for the adaptation experiment is drastically different from the community structure observed in initial metaproteomic observations when comparing acetate to WO cultures. This suggests community structure variability likely occurs during initial revival cultures from glycerol stocks prior to inoculation into experimental cultures. Meanwhile, LFQ metaproteomics revealed functional annotations that were not previously observable in acetate versus WO cultures. In particular, alkyl/aryl-sulfatase activity was found to be enriched consistently in results generated, evidenced by dodecyl sulfate metabolic process under biological processes (GO:0018909, fold change = 2.8), linear primary-alkylsulfatase activity under molecular functions (GO:0018741, fold change = 2.8), linear primary-alkylsulfatase enzyme activity (EC:3.1.6.21, fold change = 3.2), and multiple putative proteins identified via LFQ and BLAST-P annotation (fold change between 5.02 to 7.13). It is important to note that BLAST-P functional annotation is not sufficient to accurately characterize protein functionality, as sequence base alignment does not consider protein folding and structure, in contrast to more advanced tools as such AlphaFold (49), which has gained traction in studies focused on studying hypothetical proteins (50). These enzymes catalyze the hydrolytic cleavage of sulfate ester bonds in alkylbenzene sulfonates and related surfactants commonly present in petroleum products and industrial WOs (62). The degradation of alkyl sulfonates is supported by distribution of the SDS-degrading enzyme alkyl sulfatase (*SdsA1*) and its corresponding gene (sdsA1) across various strains and species of *Pseudomonas* bacteria, where *SdsA1* represents a secreted alkyl sulfatase that enables *P. aeruginosa* to utilize sodium dodecyl sulfate (SDS) as both a carbon and sulfur source (63). Furthermore, the increased protein abundance of anaerobic nitric oxide reductase transcription regulator NorR (fold change 5.59) in adapted samples shows regulation of genes involved in nitric oxide reduction, enabling bacteria to utilize alternative electron acceptors when oxygen is limited. This is found to be associated with linear alkyl sulfonate biodegradation, where increased biodegradation efficiency was reflected by enhanced nitrifying bacteria populations and upregulated ammonia monooxygenase enzyme activity, which underscores proper nutrient balancing of chemical oxygen demand to total nitrogen (COD/TN ratio) can simultaneously enhance organic pollutant degradation and nitrogen removal (64). Despite GC-MS characterization of WOs components not detecting sulfonated hydrocarbons, enzymatic and protein LFQ analysis of metaproteomes indicates the potential presence of such compounds within WOs through observed functional pathways.

ROS-AD cultures exhibited enriched acetoin catabolism regulatory protein (5.59-fold) and aconitate hydratase B (5.40-fold), which were not observed in previous metaproteome analyses. Acetoin (3-hydroxy-2-butanone) serves as an alternative carbon source for environmental bacteria when preferred substrates deplete, functioning in cellular acidification prevention and NAD+/NADH ratio maintenance (65). During acidification stress, potentially triggered by organic acid metabolites from early hydrocarbon biodegradation, bacteria convert pyruvate to acetoin via acetoin dehydrogenase, subsequently assimilating it into central carbon metabolism as acetaldehyde and acetyl-CoA (66). However, acetoin accumulation becomes toxic at high concentrations through its reactive keto group, causing DNA and protein damage (67). This toxicity correlates with enriched stress response proteins including general stress protein 18 and Protein/nucleic acid deglycase HchA (3.09-fold), which mediate protein and nucleic acid repair from aldehyde stress and methylglyoxal/glyoxal-glycated protein damage, respectively, within the DJ-1/Pfp-1 superfamily (68–70). Similar acetoin resistance mechanisms, exemplified in *Lactococcus lactis*, involve membrane stabilization through cyclopropane fatty acid modifications (67). Aconitate hydratase B (aconitase) catalyzes the stereospecific citrate-to-isocitrate conversion, serving as a key regulatory point in TCA cycle metabolism (71). Under iron-sufficient conditions, the [4Fe-4S] cluster of AcnA binds iron-responsive element (IRE) motifs, activating TCA cycle enzyme activity (72). C4-dicarboxylate transport transcriptional regulatory protein DctD (fold change 5.59) functions as the response regulator in the DctB/DctD two-component system, controlling uptake and utilization of C4-dicarboxylates (succinate, fumarate, malate, aspartate) as essential metabolic intermediates during hydrocarbon degradation (73). Collectively, these adaptations reveal a comprehensive metabolic strategy where the acetoin pathway and enhanced aconitase activity provide alternative carbon source access, and maximize energy extraction from primary substrates, while DctD-mediated C4-dicarboxylate uptake optimizes metabolic flux through central carbon pathways, enabling sustained energy production under the nutritional and environmental stresses associated with hydrocarbon exposure (74).

## Conclusion

A systematic top-down community selection strategy enriched a taxonomically diverse bacterial consortia from contaminated soil sites BPS4 and BPS8 capable of degrading complex WOs, yielding notable genera including *Pseudomonas, Achromobacter, Delftia, and Stenotrophomonas*. The strategy effectively selected microorganisms capable of utilizing petroleum hydrocarbons as sole carbon sources. Comprehensive chemical analysis revealed that the WO substrate contained volatile aromatics, polycyclic aromatic hydrocarbons (PAHs), n-alkanes, and recalcitrant organophosphates, a chemically complex environment that promoted enrichment of functionally relevant bacterial communities. We hypothesized that exposure to such chemical complexity would yield functionally versatile communities better suited for biological treatment of complex WOs. To characterize functional responses during WO biodegradation, we conducted comprehensive metaproteomic analysis on the more taxonomically diverse consortia from site BPS4. Label-free quantitative proteomics revealed taxonomic restructuring favoring *Delftia and Achromobacter* under WO sole carbon conditions, with functional enzymatic mapping demonstrating enrichment of proteins at key steps in aliphatic and aromatic degradation pathways. Consortia BPS4 preferentially routed catechol intermediates to the TCA cycle, with increased abundance of oxidoreductases, dehydrogenases, and β-oxidation enzymes providing direct molecular evidence of active hydrocarbon catabolism. Gene ontology enrichment analysis revealed concurrent adaptations including heightened nitrogen cycling, nitric oxide biosynthetic pathways, and enhanced nitrate reductase activities, functions reflecting adaptation to nutrient starvation under hydrocarbon degradation conditions. Subsequent in-vitro batch-fed adaptation of consortia BPS4 produced divergent outcomes in functional versus compositional metrics. While the community structure remained stable over multiple generations, GC-MS analysis demonstrated only limited enhancement in degrading recalcitrant oil components. However, metaproteomic analysis identified marked functional shifts: *Achromobacter* and *Pseudomonas* retained dominance while exhibiting novel alkylaryl-sulfatase activity, C4-dicarboxylate transport, acetoin catabolism, and enhanced stress response functions. These findings suggest that metabolic adaptation in complex consortia proceeds primarily through functional reorganization rather than major taxonomic restructuring, conferring improved metabolic versatility and resilience to environmental stressors.

## Acknowledgements

We thank the Defence and Security Accelerator, as part of the UK Ministry of Defence for funding this research, together with provision of environmental samples for enrichment, and waste oils. We also thank Kuwait University for funding a PhD Scholarship. The QExactive HF orbitrap mass spectrometer was funded by BBSRC UK (award no. BB/M012166/1) and major collaborations were funded by EPSRC funding (award no. EP/S020705/1).

